# Does seed mass drive interspecies variation in the effect of management practices on weed demography?

**DOI:** 10.1101/2020.01.08.898247

**Authors:** Kazakou Elena, Fried Guillaume, Cheptou Pierre-Olivier, Gimenez Olivier

## Abstract

Optimizing the effect of management practices on weed population dynamics is challenging due to the difficulties in inferring demographic parameters in seed banks and their response to disturbance. Here, we used a long-term plant survey between 2006 and 2012 in 46 French vineyards and quantified the effects of management practices (tillage, mowing and herbicide) on colonization, germination and seed survival of 30 weed species in relation to their seed mass. To do so, we used a recent statistical approach to reliably estimate demographic parameters for plant populations with a seed bank using time series of presence–absence data, which we extended to account for interspecies variation in the effects of management practices on demographic parameters. Our main finding was that when the level of disturbance increased (i.e., in plots with a higher number of herbicide, tillage or mowing treatments), colonization and survival in large-seeded species increased faster than in small-seeded species. High disturbance through tillage increased survival in the seed bank of species with high seed mass. The application of herbicides, considered as an intermediate disturbance, increased germination, survival and colonization probabilities of species with high seed mass. Mowing, representing habitats more competitive for light, increased the survival of species with high seed mass. Overall, the strong relationships between the effects of management practices and seed mass provides an indicator for predicting the dynamics of weed communities under disturbance.

## Introduction

Managing weeds is one of the most challenging issues faced by farmers as weeds can cause significant reductions in crop growth and yields especially in resource limited agroecosystems (1). In vineyards, conventional weed control methods rely on different combinations of herbicide applications, soil tillage and/or mowing applied in the vine rows and the area between them (inter-rows) (2). However, environmental concerns have brought attention to alternative weed management strategies, including the use of cover crop to improve soil organic matter content and aggregation of nutrients and water and also to control weeds (3,4). To optimize weed management, we need to not only quantify weed species presence and abundance but also to understand species demography to determine the response of weed population dynamics to disturbances.

With regard to demography, buried seeds are an essential component of the community (5) but their contribution to population dynamics is challenging to study because the seed bank is not visible. The persistence and viability of some weed species after long burial periods is well documented and seed longevity in the soil is one of the most determinant factor for the success of future generations (6). A hidden Markov model (HMM) was recently developed to estimate colonization, germination and seed bank survival using presence-absence observations (7). Here, we extend this HMM with a multilevel model in a Bayesian framework (e.g. (8)) (aka hierarchical Bayesian modeling) to determine interspecies variation in the effects of management practices on weed demography at both the plot and species level, while explicitely considering the seed bank.

Relatively undisturbed plant communities such as pasture generally exhibit low seed persistence, whereas seed persistance in frequently disturbed habitats such as arable fields will be high (9,10). Germination probability in most cases will increase after soil disturbances such as tillage and decrease after mowing (11–13). According to the disturbance intensity and frequency, germination probability can be higher in tillage systems with a frequent disturbance than in mowing systems. Additionnally, colonization can be more pronounced in frequently disturbed environments, as tillage procedures increase the seed availability and may increase seedling rates (14).

From a mechanistic perspective, we expect that seed mass, because of its influence on key processes such as dispersal in space *via* colonization and dispersal in time *via* seed bank persistence (15), will affect the influence of management practices on plant demographic parameters. Stored resources in large seeds tend to help the young seedling to survive and establish in the face of environmental hazards (e.g. deep shade, drought, herbivory). Smaller seeds also tend to be buried deeper in the soil, particularly if their shape is close to spherical, which increases their longevity in seed banks (16). Therefore, our first hypothesis is that disturbance will increase colonization and this effect will be more important for small seeds of wind-dispersed plants which should disperse at greater distances on average than large seeds (17). Our second hypothesis is that disturbance will increase germination probability and this effect will be more important for large-seeded species. Our third hypothesis is that disturbance will increase seed bank persistance and seedling of large-seeded species should better survive hazards like deep shade, physical damage and the presence of competing vegetation (18).

To explore these hypotheses, we used a unique data set covering 46 vineyard plots in France (Champagne, Beaujolais and Languedoc wine-growing areas) with 883 flora surveys performed between 2006 and 2012. First, we used our novel multilevel HMM model to test the effects of environmental variables on germination, colonization and seed persistence in the seed bank. Second, we tested whether interspecific variation in the effects of management practices (tillage, mowing and herbicide use) on demographic parameters could be explained by seed mass.

## Material and methods

### (a) The Biovigilance dataset

Vegetation surveys were conducted in French vineyards between 2006 and 2012 in the “Biovigilance” project (19), covering three main wine production regions – Languedoc, Beaujolais and northern Rhône valley and Champagne – along a latitudinal gradient of pedo-climatic conditions and management practices (for a detailed description see (20)). Languedoc has a Mediterranean climate with a mean annual temperature of 14.1 °C, and 686 mm annual rainfall in the surveyed plots (21) with a mean Treatment Frequency Index (TFI) for herbicides of 0.48, i.e. the cumulative ratio of the dose applied to the recommended dose, for all herbicide treatments applied during the growing season. Beaujolais and northern Rhone valley have a semi-continental climate with temperate influences, with a mean annual temperature of 11.4 °C and 776 mm annual rainfall with a mean TFI of 1.38. Finally, Champagne has a continental climate with oceanic influences, with a mean annual temperature of 10.1 °C and 657 mm annual rainfall with a mean TFI of 1.24.

Forty-six vineyards plots were surveyed: 18 plots in Languedoc, 18 plots in Beaujolais and northern Rhone valley and 10 plots in Champagne. In each of the 46 vineyards plot, 2,000m² quadrat surveys were performed along rows and inter-rows to account for different management practices. Following (20), we focused our analyses on the 30 most abundant species (Table S1). We extracted the presence and absence of standing weed species. Data from the presence or absence of species in the seed bank was not available and was therefore considered as a hidden variable and estimated with the probabilistic framework of HMMs (as described in the *Statistical analyses* session). We extracted seed mass values from the LEDA and TRY databases (20, 21). Seed mass, also called seed size, was defined as the oven-dry mass of a species, expressed in mg. Mean seed mass was determined by weighing the total mass of between 20 and 100 individual seeds (depending on the species), then dividing the total dry weight by the number of seeds in the sample (16). Seed mass showed high variation among species with Asteraceae species and especially *Erigeron canadensis* having the lowest value (0,0001 mg) and *Convolvulus arvensis* having the highest seed mass value (0,0145 mg

### (b) Statistical analyses

Following (7), we built a HMM by considering three states: ‘1’ for both hidden state seeds and standing flora are absent in year *t*, ‘2’ for seeds are present in year *t* but standing flora is absent in year *t*+1 and ‘3’ for seeds are present in year *t* and standing flora is present in year *t*+1, underlying the two observations made when collecting data: (i) species not seen, (ii) species seen. The HMM included four different parameters: the germination probability *g* (joint probability of seed germination success and of plant survival to adulthood), the probability of seed bank survival *s* from t to *t*+1, the probability of external colonization *c* (probability that at least one seed from outside arrived on the plot and survived to the onset of the next season) and the probability that there were seeds in the soil the year before the first observation of the existing flora in the plot *p_0_*. Parameters *g*, *s* and *c* were plot- and species-specific. Spearman rank correlation coefficients were calculated to test whether demographic parameters were correlated. First, we tested at the plot level the effects on these demographic parameters of latitude and pedoclimatic variables namely soil pH and soil texture with proportion of silt and clay that were shown to be relevant in a previous study (20). Second, we assessed the effects of management practices on the demographic parameters at the species level using species random effects on both intercepts and slopes of these relationships, and an effect of seed mass on the slope of management practices. Our multilevel HMM was fitted with a Bayesian approach (electronic supplementary material). We concluded to the significance of an effect if the 95% credible interval of the corresponding slope excluded zero. We also computed the proportion of explained variance for multilevel models (24).

## Results

### Weed demography

Seed survival probability in the seed bank *s* varied from 0.14 (*Anisantha sterilis*) to 0.82 (*Muscari neglectum*), germination probability *g* varied from 0.13 (*Chenopodium album*) to 0.79 (*Diplotaxis erucoides*) and colonization probability *c* varied from 0 (*Mercurialis annua*) to 0.86 (*Lamium amplexicaule*) (electronic supplementary material, Table S1). We found no significant correlations between the different demographic parameters (p-value>0.05).

### Effects of pedoclimatic factors on weed demography

We found no significant effect of soil parameters and latitude on colonization *c* and seed survival probability *s* (figure 1a, figure 1c). Latitude had a significant effect on germination probability *g*, with species in plots from higher latitude having a higher germination probability (figure 1b). Silt had a positive effect, albeit non-significant, on germination probability (figure 1b).

**Figure 1.**
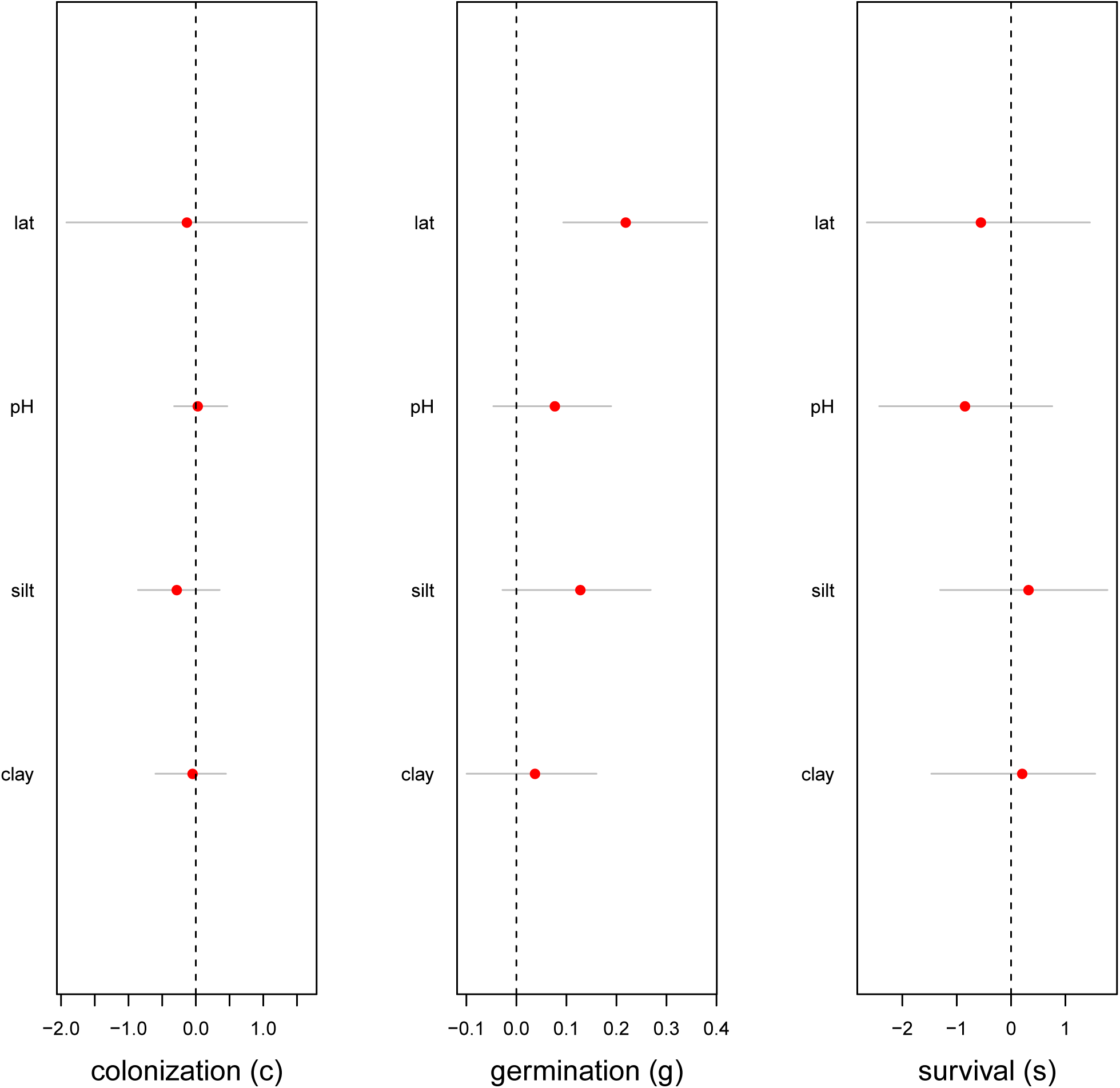
Effects of latitude and pedo-climatic variables (soil pH, proportions of silt and clay) on the demographic parameters of 30 weed species in French vineyards. Colonization (left panel), germination (middle panel) and survival (right panel) are considered. The red circles are the posterior means and the thin grey lines are the 95% credible intervals. The effect was considered significant when the corresponding credible interval did not overlap 0 represented by the dashed vertical line.

### Influence of seed mass on the effect of soil management practices on weed demography

Overall, seed mass explained interspecies variation in the effects of management practices on most of the demographic parameters (figure 2), with a positive correlation between seed mass and the intensity of these effects. Importantly, we also found that this correlation varied according to management practices.

**Figure 2.**
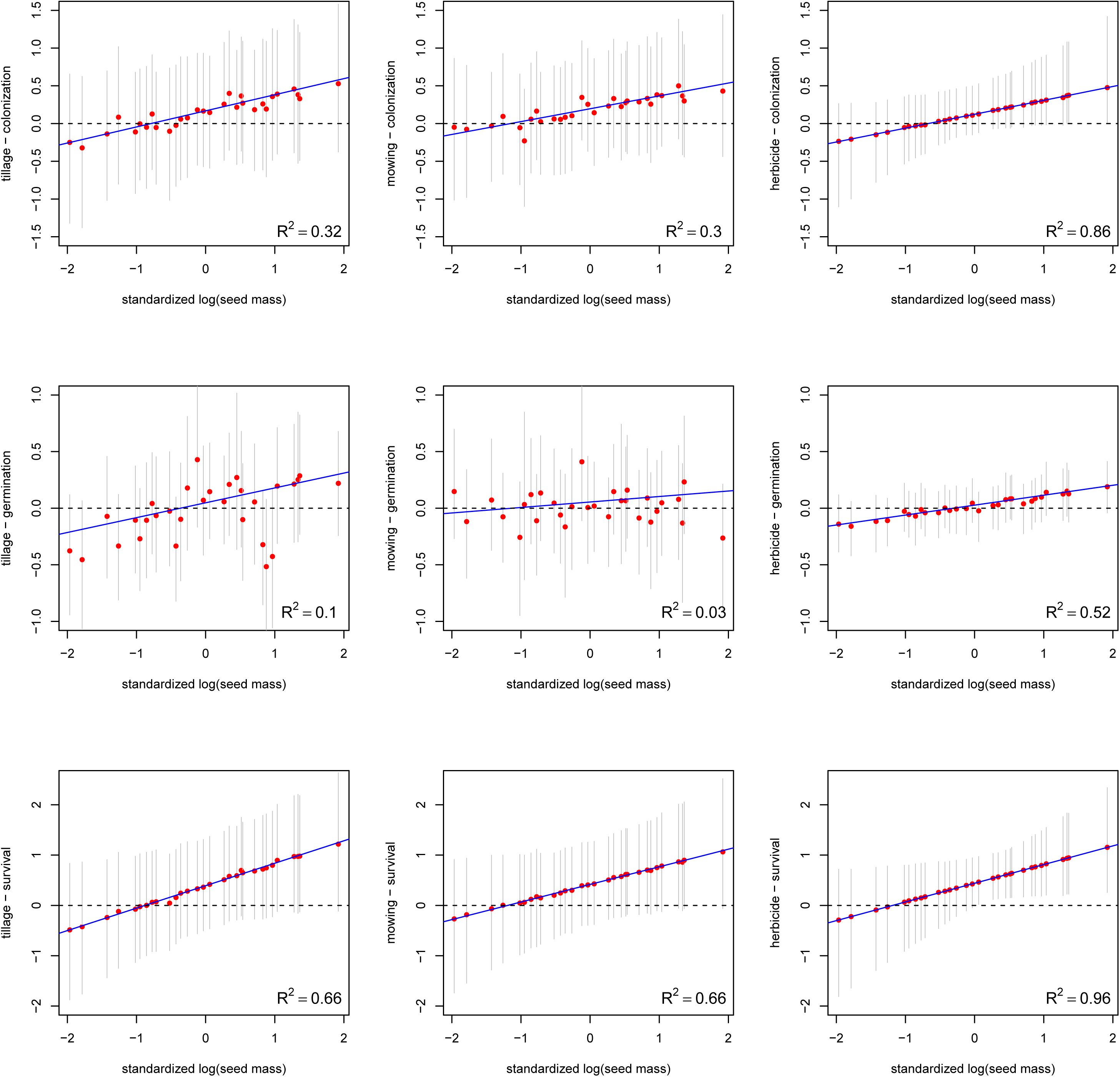
Slopes of the response of demographic parameter to management practices for 30 weed species in French vineyards. The effect of tillage (left column), mowing (middle column) and herbicides (right column) on colonization (upper row), germination (middle row) and survival (bottom row) is given as a function of seed mass on the log scale (blue solid line), as estimated by the multilevel hidden Markov model with species as a random effect, holding latitude and pedo-climatic variables to their mean values. The red circles are the posterior means and the thin grey lines are the 95% credible intervals. The proportion of explained variance is also provided (bottom right corner in each panel).

The effect of the three management practices on colonization probability covaried positively with species seed mass (figures 2a, 2b, 2c) with the strongest effet for herbicide application (R^2^ = 0.86). Herbicide application enhance more the rate of colonization for large-seeded species, like *Malva sylvestris*, *Calendula arvensis*, *Anisantha sterilis* or *Convolvulus arvensis* than small-seeded species like *Erigeron canadensis* and *Cerastium glomeratum* (figure 2c).

With regard to germination, the effect of herbicide application on vineyards covaried positively with seed mass: herbicide application increased more the germination probability of large-seeded species than that of small-seeded species (R^2^ = 0.52; figure 2f). The same result was found for the effect of tillage on germination probability (except for species *Erodium cicutarium, Fumaria officinalis, Muscari neglectum* or *Cerastium glomeratum*) although the relationship was weak (R^2^ = 0.1; figure 2d). We found no significant relationship for the effect of mowing on germination probability (figure 2e).

Finally, the effects of the three management practices on survival probability were better explained by seed mass than for germination and colonization. Seed survival in the seed bank increased more after tillage for large-seeded species (except from *Erodium cicutarium, Fumaria officinalis, Muscari neglectum*, which also had a lower increase in germination probability after the tillage) (R^2^ = 0.66; figure 2h). We found the same pattern for the effect of mowing (R^2^ = 0.66; figure 2i) and herbicide (R^2^ = 0.96; figure 2j) with species like *Malva sylvestris* or *Calendula arvensis* showing a higher increase in survival in seed bank after mowing or herbicide application than *Erigeron canadensis, Cerastium glomeratum* or *Cardamine hirsuta*.

## Discussion

### Effect of management practices on demographic parameters

Soil disturbance is one of the most important environmental filters influencing vegetation structure and composition (21, 22) and a cultivated field can be seen as the first stages of secondary succession because it is regularly disturbed. In our study, soil management practices corresponded to a gradient of disturbance frequency and intensity. Frequent mechanical weed control is expected to reduce the abundance of species with short-lived seeds and extend overall seed persistence (27). Mowing directly defoliates plants and can reduce growth, decrease plant survival and reduce or prevent seed production in two ways (28). First, mowing changes the biotic environment, such as light, temperature and soil moisture. Second, mowing changes competitive relationships between neighboring plants because different species die or regrow at different rates following mowing (29). Repeated chemical control in vineyards can be considered as an evolutionary force which results in communities secondary successions. Each year the chemical weed control restarts the succession, by the creation of gaps of bare soil but some species escape and slowly give place to a successional process (30). The positive effects of gaps in enhancing seedling recruitment are widely acknowledged (31,32).

Interestingly, our main finding was independent of the nature of farming practices and the demographic process, especially colonization and survival in seed banks. When the level of disturbance increases (i.e., in plots with a higher number of herbicide, tillage or mowing treatments), large-seeded species see colonization and survival increase more quickly than small-seeded species. We found that for large-seeded species, colonization, germination and survival in the seed bank increased with the intensity of the disturbances. This means that the intensity of the disturbances modifies the structure and community composition in the seed bank and the emerged flora. This finding can be explained by the fact that larger seeds confer an advantage with higher seedling survival and germination probability under unfavourable environment (31,33), at least on a relatively short term (34), and a greater success of emerging from burial (35,36) although they disperse less due to their larger seed mass (37). We also found that tillage increased the probability of the success of colonization from an external source and the survival to the onset of the next season for large-seeded species. One explanation is that inversion operations, like tillage, bury weed seeds at a depth where the seeds are less prone to predation and germination and thus persist longer (30). Additionnally, large-seeded species are predicted to have reduced dormancy because their seedlings can draw on a larger food reserve, and hence establish in relatively unfavorable environments. Our results showed that herbicide application increased large-seeded species germination probability and survival in seed bank. These results are in accordance with our first hypothesis that colonization should be more pronounced in frequently disturbed environments (17).

For species with very small seeds, there is not much influence of the intensity of practices. In fact, having small seeds is a typical characteristic of species with a ruderal strategy (40), which are expected to be particularly adapted to disturbed environments and capable of colonizing freshly disturbed environments (41). Our results show that for these species, the demographic parameters remain constant regardless of the level of disturbance. Their presence in the seed bank and in the emerged flora is therefore not modified by the intensity of the practices.

## Conclusion

Biodiversity conservation in vineyards is crucial not only for ensuring the sustainability and stability of farming systems, but also for providing important ecological services. However, in the last decades the vineyard has suffered an intensive management with a high mechanisation (including frequent tilling) and/or use of herbicides, which affect species richness and abundance (20). Here we used a novel statistical approach to determine the effect of management practices on seed bank dynamics in weed species. The strong relationships between these effects and seed mass provides a reliable indicator for predicting the dynamics of weed communities. A promising avenue of research is to integrate in our modeling approach biotic filters such as the relative competitive ability of weeds and the interactions between weed species.

## Author’s contributions

EK and OG conceived the study. GF oversaw data collection and compiled the dataset. OG analysed the data. EK, OG, GF and P-OC interpreted results and designed model comparisons. EK and OG framed and wrote the manuscript with input from GF and P-OC. All authors approved the final version and agree to be held accountable for the work it presents.

## Data accessibility

The data and code used in this study are available as electronic supplementary material.

