## Supplementary Material for "Does seed mass drive interspecies variation in the effect of management practices on weed demography?"

**Electronic supplementary material**

**S1**

### 1. Multilevel hidden Markov model (HMM)

To specify the HMM proposed by Pluntz et al. (2018), we need some notations first.

- Observations (or events) are $X_t = \mathrm{flora}_t$ and take values:
  - 0 is for species absent;
  - 1 is for species present.
- States are $Z_t = (\mathrm{seedbank}_{t-1}, \mathrm{flora}_t)$ and take values:
  - 1 is (0,0) for both seeds and standing flora are absent;
  - 2 is (1,0) for seeds are present but standing flora is absent on the following year;
  - 3 is (1,1) for seeds are present, and then standing flora is present.
- The parameters we need are all probabilities:
  - $g$ is the germination, i.e. the joint probability of seed germination success and of plant survival to adulthood;
  - $s$ is the seed bank survival;
  - $c$ is the external colonization, i.e. the probability that at least one seed from outside arrived on the plot and survived to the onset of the next season;
  - $p_0$ is the initialisation parameter, i.e. the probability that there were seeds in the soil the year before the first observation of the existing flora in the plot.

The HMM then consists in three components, namely the vector of initial states probabilities, the matrix of observation probabilities and the matrix of transition probabilities:

- The vector of initial probabilities is (states in columns):
$$[1-p_0 \quad p_0(1-g) \quad p_0 g];$$
- The matrix of observation probabilities is (states at $t$ in rows, observations at $t$ in columns):
$$\begin{bmatrix} 1 & 0 \\ 1 & 0 \\ 0 & 1 \end{bmatrix};$$
- The matrix of transition probabilities is (states at $t-1$ in rows, states at $t$ in columns):
$$P = \begin{bmatrix} 1-c & c(1-g) & cg \\ (1-c)(1-s) & (1-g)(1-(1-c)(1-s)) & g(1-(1-c)(1-s)) \\ 0 & 1-g & g \end{bmatrix}.$$

Last, we consider the effect of covariates on the demographic parameters $g$, $s$ and $c$ which we denote $\theta$ for short. We write $i$ for the plot index and $j$ for the species index:

$$\mathrm{logit}(\theta(i,j)) = \alpha_0(j) + \alpha_1(j)\ \mathrm{latitude}(i) + \alpha_2(j)\ \mathrm{pH}(i) + \alpha_3(j)\ \mathrm{silt}(i) + \alpha_4(j)\ \mathrm{clay}(i)$$
$$+\alpha_5(j)\ \mathrm{mowing}(i) + \alpha_6(j)\ \mathrm{tillage}(i) + \alpha_7(j)\ \mathrm{herbicide}(i)$$

with the following random effects:

$$\alpha_k(j) \sim N(\overline{\alpha}_k, \sigma_k^2), k = 0,1,2,3,4$$

and

$$\alpha_k(j) \sim N(\gamma_k + \beta_k \mathrm{seedmass}(j), \sigma_k^2), k = 5,6,7.$$

#### 2. Run the analysis

To implement the Bayesian analysis of our multilevel HMM, we used non-informative normal prior distributions for the regression coefficients and uniform prior distributions for the standard deviation of the random effects. We ran two MCMC in parallel with different initial values, 10,000 iterations each and an initial burn-in of 2,500 iterations. We assessed convergence by visual inspection and by using the Gelman and Rubin R-hat diagnostic. We used OpenBUGS which allows parallel computation.

First, load the packages we need:

```
library(R2OpenBUGS) # bayesian analyses
library(coda) # bayesian diagnostics
library(mcmcplots) # viz mcmc
library(snow) # parallelization
library(snowfall) # parallelization
library(tidyverse) # data viz and manipulation
```

```
## ── Attaching packages ──────────────────────────────────────────────
──────────── tidyverse 1.2.1 ──

## ✔ ggplot2 3.2.0          ✔ purrr   0.3.2.9000
## ✔ tibble  2.1.3          ✔ dplyr   0.8.3
## ✔ tidyr   1.0.0          ✔ stringr 1.4.0
## ✔ readr   1.3.1          ✔ forcats 0.4.0

## ── Conflicts ────────────────────────────────────────────────────────
────── tidyverse_conflicts() ──
## ✖ dplyr::filter() masks stats::filter()
## ✖ dplyr::lag()    masks stats::lag()
```

We run two chains in parallel. To do so, we set the number of CPUs to two and assign the R2OpenBUGS library to each CPU:

```
sfInit(parallel=TRUE, cpus=2)

## Warning in searchCommandline(parallel, cpus = cpus, type = type,
## socketHosts = socketHosts, : Unknown option on commandline:
## rmarkdown::render('/Users/oliviergimenez/Desktop/appendix/
## appendix.Rmd',~+~~+~encoding~+~

## R Version:  R version 3.6.1 (2019-07-05)

## snowfall 1.84-6.1 initialized (using snow 0.4-3): parallel execution o
n 2 CPUs.

sfLibrary(R2OpenBUGS)

## Library R2OpenBUGS loaded.

## Library R2OpenBUGS loaded in cluster.
```

Now get the data and format them for analysis in OpenBUGS:

```
load('dat.RData')
mydatax <- list(nsp = nsp, # number of species
                nsites = nsites, # number of plots
                nocc = nocc, # number of sampling occasions
                mydata = sp_histories_array, # detections/non-detections
                chim = chim, # herbicide
                mec = mec, # tillage
                fau = fau, # mowin
                lat = lat, # latitude
                seedmass = seedmass, # species seed mass
                ph = ph, # pH
                silt = silt, # % silt
                clay = clay) # % clay
```

Create separate directory for each CPU process:

```
folder1 <- paste(getwd(), "/chain1", sep="")
folder2 <- paste(getwd(), "/chain2", sep="")
##### dir.create(folder1); dir.create(folder2); # uncomment if  you'd lik
e to run the analysis
                                                # (warning: takes several
hours)
```

Now specify the multilevel HMM model:

```
# sinking the model into a file in each directory
for (folder in c(folder1, folder2)){
  sink(paste(folder, "/nummodel.txt", sep=""))
cat("
    model{
  # DEFINE PARAMETERS

  # OBSERVATION PROCESS: probabilities of observations (columns) at a giv
en occasion
  #                      given states (rows) at this occasion
  po[1,1] <- 1
  po[1,2] <- 0
  po[2,1] <- 1
  po[2,2] <- 0
  po[3,1] <- 0
  po[3,2] <- 1

  po.init[1,1] <- 1
  po.init[1,2] <- 0
  po.init[2,1] <- 1
  po.init[2,2] <- 0
  po.init[3,1] <- 0
  po.init[3,2] <- 1

  # STATE PROCESS: probabilities of states at t+1 (columns) given states
at t (rows)
```

```
# probabilities for each INITIAL STATES
for (s in 1:nsp){ # for each species
  for (i in 1:nsites){ # for each site
    px0[s,i,1] <- 1 - p_knot[s]
    px0[s,i,2] <- p_knot[s] * (1 - gg[s,i])
    px0[s,i,3] <- p_knot[s] * gg[s,i]
    px[s,i,1,1] <- 1 - cc[s,i]
    px[s,i,1,2] <- (1 - gg[s,i]) * cc[s,i]
    px[s,i,1,3] <- gg[s,i] * cc[s,i]
    px[s,i,2,1] <- (1 - cc[s,i]) * (1 - ss[s,i])
    px[s,i,2,2] <- (1 - gg[s,i]) * (1 - (1 - cc[s,i]) * (1 - ss[s,i]))
    px[s,i,2,3] <- gg[s,i] * (1 - (1 - cc[s,i]) * (1 - ss[s,i]))
    px[s,i,3,1] <- 0
    px[s,i,3,2] <- 1 - gg[s,i]
    px[s,i,3,3] <- gg[s,i]
  } # end site
} # end species

# LIKELIHOOD

for (s in 1:nsp){ # for each species
  for (i in 1:nsites){ # for each ind within species
    # estimated probabilities of initial states are the proportions
    # in each state at first capture occasion
    alive[s,i,1] ~ dcat(px0[s,i,1:3])
    mydata[s,i,1] ~ dcat(po.init[alive[s,i,1],1:2])
    for (j in 2:nocc){  # loop over time

      ## STATE EQUATIONS ##
      # draw states at j given states at j-1
      alive[s,i,j] ~ dcat(px[s,i,alive[s,i,j-1],1:3])

      ## OBSERVATION EQUATIONS ##
      # draw observations at j given states at j
      mydata[s,i,j] ~ dcat(po[alive[s,i,j],1:2])

    } # end time
  } # end ind
} # end species

# PRIORS
for (s in 1:nsp){ # for each species
  logit(p_knot[s]) <- intpknot + epspknot[s]
  epspknot[s] ~ dnorm(0, tauknot)
  for (i in 1:nsites){ # for each ind within species
  logit(cc[s,i]) <- intercept[1,s] +
    slopelat[1,s] * lat[i] +
    slopeph[1,s] * ph[i] +
    slopesilt[1,s] * silt[i] +
    slopeclay[1,s] * clay[i] +
    slopemeca[1,s] * mec[i] +
```

```
    slopechim[1,s] * chim[i] +
    slopefau[1,s] * fau[i]
  logit(gg[s,i]) <- intercept[2,s] +
    slopelat[2,s] * lat[i] +
    slopeph[2,s] * ph[i] +
    slopesilt[2,s] * silt[i] +
    slopeclay[2,s] * clay[i] +
    slopemeca[2,s] * mec[i] +
    slopechim[2,s] * chim[i] +
    slopefau[2,s] * fau[i]
  logit(ss[s,i]) <- intercept[3,s] +
    slopelat[3,s] * lat[i] +
    slopeph[3,s] * ph[i] +
    slopesilt[3,s] * silt[i] +
    slopeclay[3,s] * clay[i] +
    slopemeca[3,s] * mec[i] +
    slopechim[3,s] * chim[i] +
    slopefau[3,s] * fau[i]
  }
}

intpknot ~ dnorm(0, 1)
tauknot <- 1 / (sdpknot * sdpknot)
sdpknot ~ dunif(0, 10)

  for (i in 1:3){ # for each parameter: cc, gg and ss

    meanint[i] ~ dnorm(0, 1)
    tauint[i] <- 1 / (sdint[i] * sdint[i])
    sdint[i] ~ dunif(0, 10)

    meanlat[i] ~ dnorm(0, 1)
    taulat[i] <- 1 / (sdlat[i] * sdlat[i])
    sdlat[i] ~ dunif(0, 10)

    meanph[i] ~ dnorm(0, 1)
    tauph[i] <- 1 / (sdph[i] * sdph[i])
    sdph[i] ~ dunif(0, 10)

    meansilt[i] ~ dnorm(0, 1)
    tausilt[i] <- 1 / (sdsilt[i] * sdsilt[i])
    sdsilt[i] ~ dunif(0, 10)

    meanclay[i] ~ dnorm(0, 1)
    tauclay[i] <- 1 / (sdclay[i] * sdclay[i])
    sdclay[i] ~ dunif(0, 10)

    ameca[i] ~ dunif(0, 1)
    bmeca[i] ~ dunif(0, 1)
    achim[i] ~ dunif(0, 1)
    bchim[i] ~ dunif(0, 1)
    afau[i] ~ dunif(0, 1)
```

```
      bfau[i] ~ dunif(0, 1)

      for (s in 1:nsp){ # for each species
        intercept[i,s] ~ dnorm(meanint[i], tauint[i])
        slopelat[i,s] ~ dnorm(meanlat[i], taulat[i])
        slopeph[i,s] ~ dnorm(meanph[i], tauph[i])
        slopesilt[i,s] ~ dnorm(meansilt[i], tausilt[i])
        slopeclay[i,s] ~ dnorm(meanclay[i], tauclay[i])

        slopemeca[i,s] ~ dnorm(slopemeca1[i,s], taumeca)
        slopemeca1[i,s] <- ameca[i] + bmeca[i] * seedmass[s]
        e.meca[i,s] <- slopemeca[i,s] - slopemeca1[i,s]

        slopechim[i,s] ~ dnorm(slopechim1[i,s], tauchim)
        slopechim1[i,s] <- achim[i] + bchim[i] * seedmass[s]
        e.chim[i,s] <- slopechim[i,s] - slopechim1[i,s]

        slopefau[i,s] ~ dnorm(slopefau1[i,s], taufau)
        slopefau1[i,s] <- afau[i] + bfau[i] * seedmass[s]
        e.fau[i,s] <- slopefau[i,s] - slopefau1[i,s]

    }
  }

taumeca <- 1 / (sdmeca * sdmeca)
sdmeca ~ dunif(0, 10)
tauchim <- 1 / (sdchim * sdchim)
sdchim ~ dunif(0, 10)
taufau <- 1 / (sdfau * sdfau)
sdfau ~ dunif(0, 10)

}
")
sink()
}
```

Define the function that will run MCMC on each CPU:

```
# Arguments:
# chain - will be 1 or 2
# x.data - the data list
# params - parameters to be monitored
parallel.bugs <- function(chain, x.data, params)
{
  # a. defining directory for each CPU
  sub.folder <- paste(getwd(),"/chain", chain, sep="")

  # b. specifying the initial MCMC values
  inits <- function()list(meanint = rnorm(3, 0, 1),
                          meanlat = rnorm(3, 0, 1),
                          meanph = rnorm(3, 0, 1),
                          meansilt = rnorm(3, 0, 1),
                          meanclay = rnorm(3, 0, 1))
```

```
  # c. calling OpenBugs
  bugs(data= x.data, inits=inits, parameters.to.save= params,
             n.iter = 10000, n.chains=1,
             model.file="nummodel.txt", debug=T, codaPkg=TRUE,
             useWINE=TRUE,
             OpenBUGS.pgm = "/Applications/OpenBUGS323/OpenBUGS.exe",
             working.directory = sub.folder,
             WINE="/usr/local/Cellar/wine/2.0.4/bin/wine",
             WINEPATH= "/usr/local/Cellar/wine/2.0.4/bin/winepath")
}
```

Then specify the parameters to be monitored:

```
parameters <- c("p_knot","intpknot","sdpknot","meanint","meanlat","meanph
","meansilt",
                "meanclay","ameca","bmeca","achim","bchim","afau","bfau",
"intercept",
                "slopelat","slopeph","slopesilt","slopeclay","slopemeca",
"slopechim",
                "slopefau","sdint","sdlat","sdph","sdsilt","sdclay","sdme
ca","sdchim",
                "sdfau","e.meca","e.chim","e.fau")
```

Now the code to fit the model. Note that we do not run it as the analysis takes several hours to complete.

```
# calling the sfLapply function that will run
# parallel.bugs on each of the 2 CPUs
start_time <- Sys.time()
sfLapply(1:2, fun=parallel.bugs, x.data=mydatax, params=parameters)
end_time <- Sys.time()
end_time - start_time #
```

For convenience, we provide the results and post-process them. You may download the MCMC outputs from *https://mycore.core-cloud.net/index.php/s/YUDN93cjeOugozH*, then load them in R:

```
load('MCMCoutputs.Rdata')
```

If you'd like to do it yourself, you'll need the following lines of codes:

```
# locating position of each CODA chain and read them
chain1 <- paste(folder1, "/CODAchain1.txt", sep="")
chain2 <- paste(folder2, "/CODAchain1.txt", sep="")
res <- read.bugs(c(chain1, chain2), quiet=TRUE) # takes a minute or so
out2 <- as.mcmc(rbind(res[[1]],res[[2]]))
save(out2, file="MCMCoutputs.RData", compress="xz")
```

Rename relevant variables:

```
varnames(out2)[1:18] <- c("chim_cc","chim_gg","chim_ss","fau_cc","fau_gg"
,"fau_ss","meca_cc",
                          "meca_gg","meca_ss","slope_cc_chim","slope_gg_c
him","slope_ss_chim",
```

```
                          "slope_cc_fau","slope_gg_fau","slope_ss_fau","s
lope_cc_meca",
                          "slope_gg_meca","slope_ss_meca")
varnames(out2)[381:383] <- c("clay_cc","clay_gg","clay_ss")
varnames(out2)[387:395] <- c("lat_cc","lat_gg","lat_ss","ph_cc","ph_gg","
ph_ss","silt_cc",
                             "silt_gg","silt_ss")
```

Get Figure 1:

```
posterior.medians <- apply(out2, 2, median)
#pdf('fig1.pdf')
par(mfrow=c(1,3))
caterplot(out2,
          parms = c("lat_cc","ph_cc","silt_cc","clay_cc"),
          collapse=FALSE,
          reorder = FALSE,
          style = 'plain',
          cex.labels = 1,
          labels = c( 'lat', 'pH', 'silt', 'clay'),
          quantiles = list(outer=c(0.025,0.975),
                           inner=c(0.025,0.975)),
          col='gray',
          lwd=1)
caterpoints(posterior.medians[c("lat_cc","ph_cc","silt_cc","clay_cc")],
            pch=19,
            col="red")
abline(v = 0,
       col = 'black',
       lty = 2)
mtext('colonization (c)',
      side = 1,
      line = 3,
      cex = 1)
caterplot(out2,
          parms = c("lat_gg","ph_gg","silt_gg","clay_gg"),
          collapse=FALSE,
          reorder = FALSE,
          style = 'plain',
          cex.labels = 1,
          labels = c( 'lat', 'pH', 'silt', 'clay'),
          quantiles = list(outer=c(0.025,0.975),
                           inner=c(0.025,0.975)),
          col='gray',
          lwd=1)
caterpoints(posterior.medians[c("lat_gg","ph_gg","silt_gg","clay_gg")],
            pch=19,
            col="red")
abline(v = 0,
       col = 'black',
       lty = 2)
mtext('germination (g)',
```

```
      side = 1,
      line = 3,
      cex = 1)
caterplot(out2,
          parms = c("lat_ss","ph_ss","silt_ss","clay_ss"),
          collapse=FALSE,
          reorder = FALSE,
          style = 'plain',
          cex.labels = 1,
          labels = c( 'lat', 'pH', 'silt', 'clay'),
          quantiles = list(outer=c(0.025,0.975),
                           inner=c(0.025,0.975)),
          col='gray',
          lwd=1)
caterpoints(posterior.medians[c("lat_ss","ph_ss","silt_ss","clay_ss")],
            pch=19,
            col="red")
abline(v = 0,
       col = 'black',
       lty = 2)
mtext('survival (s)',
      side = 1,
      line = 3,
      cex = 1)
```

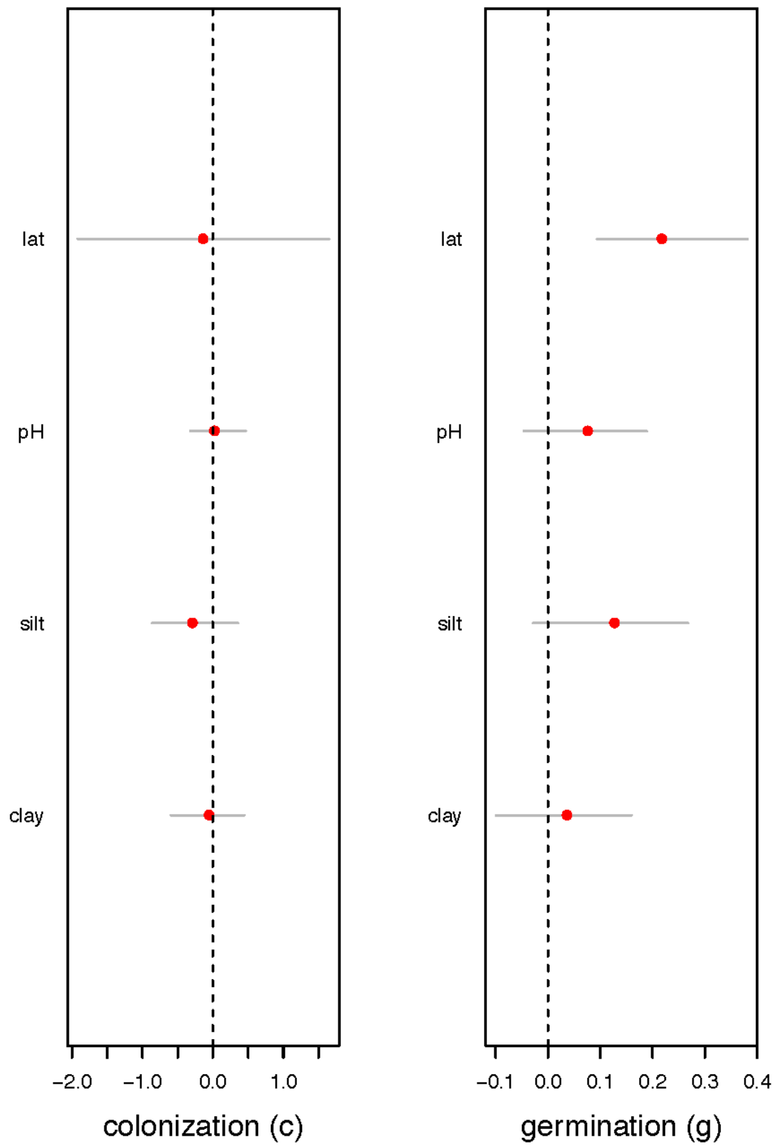


```
#dev.off()
```

Get proportion of explained variance:

```
names_col <- colnames(out2)

# R2 for meca
e.prac <- out2[,grep('e.meca',names_col)]
eprac <- vector("list", 3) # col, ger, sur
eprac[[1]] <- e.prac[,1:30]
eprac[[2]] <- e.prac[,31:60]
eprac[[3]] <- e.prac[,61:90]
slope <- out2[,grep('slopemeca',names_col)]
b <- vector("list", 3)
b[[1]] <- slope[,1:30]
b[[2]] <- slope[,31:60]
b[[3]] <- slope[,61:90]
```

```
rsquared.meca <- rep(NA,3)
#lambda.meca <- rep(NA,3)

for (i in 1:3){
  rsquared.meca[i] <- 1 - mean(apply (eprac[[i]], 1, var)) / mean (apply
(b[[i]], 1, var))
}

# R2 for chim
e.prac <- out2[,grep('e.chim',names_col)]
eprac <- vector("list", 3) # col, ger, sur
eprac[[1]] <- e.prac[,1:30]
eprac[[2]] <- e.prac[,31:60]
eprac[[3]] <- e.prac[,61:90]
slope <- out2[,grep('slopechim',names_col)]
b <- vector("list", 3)
b[[1]] <- slope[,1:30]
b[[2]] <- slope[,31:60]
b[[3]] <- slope[,61:90]

rsquared.chim <- rep(NA,3)
#lambda.meca <- rep(NA,3)

for (i in 1:3){
  rsquared.chim[i] <- 1 - mean(apply (eprac[[i]], 1, var)) / mean (apply
(b[[i]], 1, var))
}


# R2 for fau
e.prac <- out2[,grep('e.fau',names_col)]
eprac <- vector("list", 3) # col, ger, sur
eprac[[1]] <- e.prac[,1:30]
eprac[[2]] <- e.prac[,31:60]
eprac[[3]] <- e.prac[,61:90]
slope <- out2[,grep('slopefau',names_col)]
b <- vector("list", 3)
b[[1]] <- slope[,1:30]
b[[2]] <- slope[,31:60]
b[[3]] <- slope[,61:90]

rsquared.fau <- rep(NA,3)
#lambda.meca <- rep(NA,3)

for (i in 1:3){
  rsquared.fau[i] <- 1 - mean(apply (eprac[[i]], 1, var)) / mean (apply (
b[[i]], 1, var))
}

rsquared.meca # R2 for tillage and 3 demographic parameters

## [1] 0.3197832 0.1028678 0.6617302
```

```
rsquared.chim # R2 for herbicide and 3 demographic parameters

## [1] 0.8584530 0.5220513 0.9600250

rsquared.fau # R2 for mowing and 3 demographic parameters

## [1]  0.29789503 -0.03217493  0.65964770
```

Last, get Figure 2:

```
species <- c('CONAR','CIRAR','SENVU','DIPER','GERRT','ERICA','TAROF','CVP
SA','LACSE','SONOL',
             'VERPE','POAAN','EROCI','CHEAL','PLALA','STEME','MALSI','DAU
CA','GERCO','FUMOF',
             'CARHI','PICEC','SONAS','MERAN','BROST','CERGL','LOLMU','MUS
RA','LAMAM','CLDAR')
names <- species %>%
  enframe() %>%
  mutate(species = tolower(species)) %>%
  pull(species)
seedmassobs <- seedmass
ord <- order(seedmassobs)
names[ord]

##  [1] "erica" "cergl" "carhi" "cvpsa" "diper" "senvu" "sonas" "sonol"
##  [9] "poaan" "steme" "lacse" "lamam" "cheal" "tarof" "picec" "verpe"
## [17] "cirar" "plala" "meran" "gerrt" "dauca" "lolmu" "eroci" "fumof"
## [25] "musra" "gerco" "malsi" "cldar" "brost" "conar"

#pdf('fig2.pdf', width = 10, # The width of the plot in inches
#    height = 10) # The height of the plot in inches

par(mfrow=c(3,3))
# cc and meca
fac <- out2[,grep("slopemeca\\[1,", varnames(out2))]
#fac <- rbind(fac[[1]],fac[[2]])
mean_species <- apply(fac,2,mean)
q25 <- apply(fac,2,quantile, probs = 2.5/100)
q975 <- apply(fac,2,quantile, probs = 97.5/100)
plot(seedmassobs[ord],
     mean_species[ord],
     type='n',
     ylim=c(-1.5,1.5),
xlab='standardized log(seed mass)',ylab='tillage - colonization',xaxt="n"
)
axis(1)
for (i in 1:30){
    segments(seedmassobs[ord][i],q25[ord][i],seedmassobs[ord][i],q975[ord
][i],
             col='grey',pch=19,cex=0.5,lwd=0.6)
    points(seedmassobs[ord][i],mean_species[ord][i],col='red',cex=0.7, pc
h=19)
}
```

```
abline(c(mean(unlist(out2[,'meca_cc'])), mean(unlist(out2[,'slope_cc_meca
']))), col = 'blue')
abline(h = 0, lty = 2)
text(1.5, -1.4, expression(R^2 == 0.32), cex = 1.2)

# cc and fau
fac <- out2[,grep("slopefau\\[1,", varnames(out2))]
#fac <- rbind(fac[[1]],fac[[2]])
mean_species <- apply(fac,2,mean)
q25 <- apply(fac,2,quantile, probs = 2.5/100)
q975 <- apply(fac,2,quantile, probs = 97.5/100)
plot(seedmassobs[ord],
     mean_species[ord],
     type='n',
     ylim=c(-1.5,1.5),
xlab='standardized log(seed mass)',ylab='mowing - colonization',xaxt="n")
axis(1)
for (i in 1:30){
    segments(seedmassobs[ord][i],q25[ord][i],seedmassobs[ord][i],q975[ord
][i],
             col='grey',pch=19,cex=0.5,lwd=0.6)
    points(seedmassobs[ord][i],mean_species[ord][i],col='red',cex=0.7, pc
h=19)
}
abline(c(mean(unlist(out2[,'fau_cc'])), mean(unlist(out2[,'slope_cc_fau']
))), col = 'blue')
abline(h = 0, lty = 2)
text(1.5, -1.4, expression(R^2 == 0.30), cex = 1.2)

# cc and chimie
fac <- out2[,grep("slopechim\\[1,", varnames(out2))]
#fac <- rbind(fac[[1]],fac[[2]])
mean_species <- apply(fac,2,mean)
q25 <- apply(fac,2,quantile, probs = 2.5/100)
q975 <- apply(fac,2,quantile, probs = 97.5/100)
plot(seedmassobs[ord],
     mean_species[ord],
     type='n',
     ylim=c(-1.5,1.5),
xlab='standardized log(seed mass)',ylab='herbicide - colonization',xaxt="
n")
axis(1)
for (i in 1:30){
    segments(seedmassobs[ord][i],q25[ord][i],seedmassobs[ord][i],q975[ord
][i],
             col='grey',pch=19,cex=0.5,lwd=0.6)
    points(seedmassobs[ord][i],mean_species[ord][i],col='red',cex=0.7, pc
h=19)
}
abline(c(mean(unlist(out2[,'chim_cc'])), mean(unlist(out2[,'slope_cc_chim
']))), col = 'blue')
abline(h = 0, lty = 2)
```

```
text(1.5, -1.4, expression(R^2 == 0.86), cex = 1.2)

# gg and meca
fac <- out2[,grep("slopemeca\\[2,", varnames(out2))]
#fac <- rbind(fac[[1]],fac[[2]])
mean_species <- apply(fac,2,mean)
q25 <- apply(fac,2,quantile, probs = 2.5/100)
q975 <- apply(fac,2,quantile, probs = 97.5/100)
plot(seedmassobs[ord],
     mean_species[ord],
     type='n',
     ylim=c(-1,1),
xlab='standardized log(seed mass)',ylab='tillage - germination',xaxt="n")
axis(1)
for (i in 1:30){
    segments(seedmassobs[ord][i],q25[ord][i],seedmassobs[ord][i],q975[ord
][i],
             col='grey',pch=19,cex=0.5,lwd=0.6)
    points(seedmassobs[ord][i],mean_species[ord][i],col='red',cex=0.7, pc
h=19)
}
abline(c(mean(unlist(out2[,'meca_gg'])), mean(unlist(out2[,'slope_gg_meca
']))), col = 'blue')
abline(h = 0, lty = 2)
text(1.5, -0.9, expression(R^2 == 0.10), cex = 1.2)

# gg and fau
fac <- out2[,grep("slopefau\\[2,", varnames(out2))]
#fac <- rbind(fac[[1]],fac[[2]])
mean_species <- apply(fac,2,mean)
q25 <- apply(fac,2,quantile, probs = 2.5/100)
q975 <- apply(fac,2,quantile, probs = 97.5/100)
plot(seedmassobs[ord],
     mean_species[ord],
     type='n',
     ylim=c(-1,1),
xlab='standardized log(seed mass)',ylab='mowing - germination',xaxt="n")
axis(1)
for (i in 1:30){
    segments(seedmassobs[ord][i],q25[ord][i],seedmassobs[ord][i],q975[ord
][i],
             col='grey',pch=19,cex=0.5,lwd=0.6)
    points(seedmassobs[ord][i],mean_species[ord][i],col='red',cex=0.7, pc
h=19)
}
abline(c(mean(unlist(out2[,'fau_gg'])), mean(unlist(out2[,'slope_gg_fau']
))), col = 'blue')
abline(h = 0, lty = 2)
text(1.5, -0.9, expression(R^2 == 0.03), cex = 1.2)

# gg and chimie
fac <- out2[,grep("slopechim\\[2,", varnames(out2))]
```

```
#fac <- rbind(fac[[1]],fac[[2]])
mean_species <- apply(fac,2,mean)
q25 <- apply(fac,2,quantile, probs = 2.5/100)
q975 <- apply(fac,2,quantile, probs = 97.5/100)
plot(seedmassobs[ord],
     mean_species[ord],
     type='n',
     ylim=c(-1,1),
xlab='standardized log(seed mass)',ylab='herbicide - germination',xaxt="n
")
axis(1)
for (i in 1:30){
    segments(seedmassobs[ord][i],q25[ord][i],seedmassobs[ord][i],q975[ord
][i],
             col='grey',pch=19,cex=0.5,lwd=0.6)
    points(seedmassobs[ord][i],mean_species[ord][i],col='red',cex=0.7, pc
h=19)
}
abline(c(mean(unlist(out2[,'chim_gg'])), mean(unlist(out2[,'slope_gg_chim
']))), col = 'blue')
abline(h = 0, lty = 2)
text(1.5, -0.9, expression(R^2 == 0.52), cex = 1.2)

# ss and meca
fac <- out2[,grep("slopemeca\\[3,", varnames(out2))]
#fac <- rbind(fac[[1]],fac[[2]])
mean_species <- apply(fac,2,mean)
q25 <- apply(fac,2,quantile, probs = 2.5/100)
q975 <- apply(fac,2,quantile, probs = 97.5/100)
plot(seedmassobs[ord],
     mean_species[ord],
     type='n',
     ylim=c(-2,2.5),
xlab='standardized log(seed mass)',ylab='tillage - survival',xaxt="n")
axis(1)
for (i in 1:30){
    segments(seedmassobs[ord][i],q25[ord][i],seedmassobs[ord][i],q975[ord
][i],
             col='grey',pch=19,cex=0.5,lwd=0.6)
    points(seedmassobs[ord][i],mean_species[ord][i],col='red',cex=0.7, pc
h=19)
}
abline(c(mean(unlist(out2[,'meca_ss'])), mean(unlist(out2[,'slope_ss_meca
']))), col = 'blue')
abline(h = 0, lty = 2)
text(1.5, -1.8, expression(R^2 == 0.66), cex = 1.2)

# ss and fau
fac <- out2[,grep("slopefau\\[3,", varnames(out2))]
#fac <- rbind(fac[[1]],fac[[2]])
mean_species <- apply(fac,2,mean)
q25 <- apply(fac,2,quantile, probs = 2.5/100)
```

```
q975 <- apply(fac,2,quantile, probs = 97.5/100)
plot(seedmassobs[ord],
     mean_species[ord],
     type='n',
     ylim=c(-2,2.5),
xlab='standardized log(seed mass)',ylab='mowing - survival',xaxt="n")
axis(1)
for (i in 1:30){
    segments(seedmassobs[ord][i],q25[ord][i],seedmassobs[ord][i],q975[ord
][i],
             col='grey',pch=19,cex=0.5,lwd=0.6)
    points(seedmassobs[ord][i],mean_species[ord][i],col='red',cex=0.7, pc
h=19)
}
abline(c(mean(unlist(out2[,'fau_ss'])), mean(unlist(out2[,'slope_ss_fau']
))), col = 'blue')
abline(h = 0, lty = 2)
text(1.5, -1.8, expression(R^2 == 0.66), cex = 1.2)

# ss and chimie
fac <- out2[,grep("slopechim\\[3,", varnames(out2))]
#fac <- rbind(fac[[1]],fac[[2]])
mean_species <- apply(fac,2,mean)
q25 <- apply(fac,2,quantile, probs = 2.5/100)
q975 <- apply(fac,2,quantile, probs = 97.5/100)
plot(seedmassobs[ord],
     mean_species[ord],
     type='n',
     ylim=c(-2,2.5),
xlab='standardized log(seed mass)',ylab='herbicide - survival',xaxt="n")
axis(1)
for (i in 1:30){
    segments(seedmassobs[ord][i],q25[ord][i],seedmassobs[ord][i],q975[ord
][i],
             col='grey',pch=19,cex=0.5,lwd=0.6)
    points(seedmassobs[ord][i],mean_species[ord][i],col='red',cex=0.7, pc
h=19)
}
abline(c(mean(unlist(out2[,'chim_ss'])), mean(unlist(out2[,'slope_ss_chim
']))), col = 'blue')
abline(h = 0, lty = 2)
text(1.5, -1.8, expression(R^2 == 0.96), cex = 1.2)
```

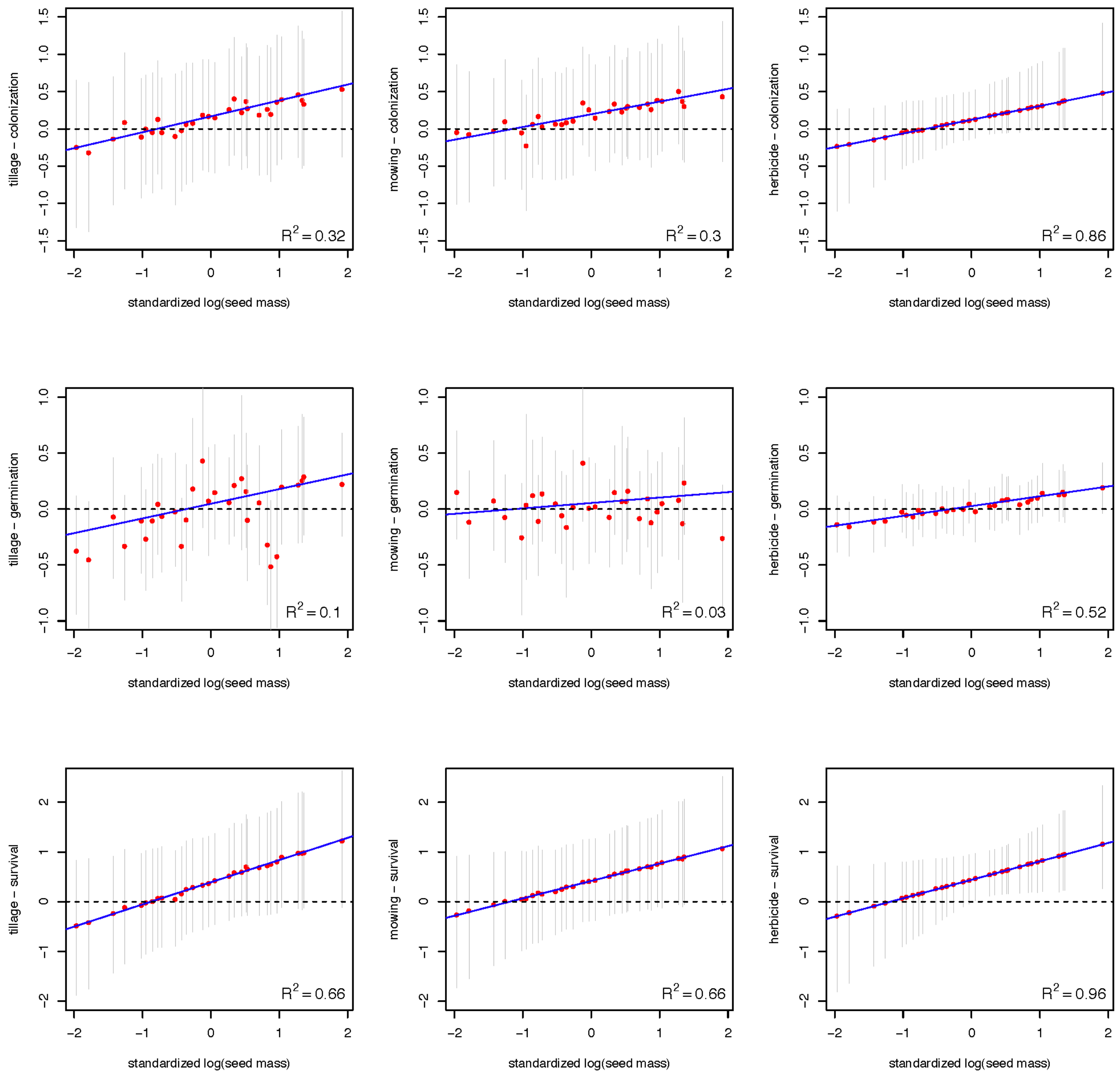


```
#dev.off()
```

We provide below the 95% credible intervals for the slopes of the relationships in Figure 2 (slope of the blue solid lines):

beta c-tillage: (0.01, 0.64)
beta c-mowing: (0.01, 0.54)
beta c-herbicide: (0.00, 0.61)
beta g-tillage: (0.01, 0.31)
beta g-mowing: (0.00, 0.16)
beta g-herbicide: (0.01, 0.20)
beta s-tillage: (0.02, 0.96)
beta s-mowing: (0.01, 0.95)
beta s-herbicide: (0.02, 0.98).

Also, here is the range of values for the three treatments:

- herbicide:

| Min. | 1st Qu. | Median | Mean | 3rd Qu. | Max. |
|---|---|---|---|---|---|
| 0.00 | 0.00 | 1.00 | 1.75 | 3.00 | 7.00 |

- tillage:

| Min. | 1st Qu. | Median | Mean | 3rd Qu. | Max. |
|---|---|---|---|---|---|
| 0.000 | 1.000 | 3.500 | 4.705 | 8.000 | 18.000 |

- mowing:

| Min. | 1st Qu. | Median | Mean | 3rd Qu. | Max. |
|---|---|---|---|---|---|
| 0.00 | 0.00 | 0.50 | 2.75 | 4.00 | 14.00 |

**Table S1:** Parameter estimates from the Biovigilance dataset. Posterior means are provided. Values for mean seed mass from LEDA and TRY database are given.

| Species name | Colonization | Germination | Survival | Seed mass (g) |
|---|---|---|---|---|
| *Anisantha sterilis* | 0,01 | 0,24 | 0,14 | 0,0064 |
| *Calendula arvensis* | 0,13 | 0,17 | 0,68 | 0,0001 |
| *Cardamine hirsuta* | 0,54 | 0,2 | 0,48 | 0,0001 |
| *Cerastium glomeratum* | 0,05 | 0,41 | 0,22 | 0,0006 |
| *Chenopodium album* | 0,19 | 0,13 | 0,46 | 0,0013 |
| *Cirsium arvense* | 0,74 | 0,41 | 0,6 | 0,0062 |
| *Convolvulus arvensis* | 0,69 | 0,19 | 0,45 | 0,0145 |
| *Crepis sancta* | 0,52 | 0,56 | 0,69 | 0,0001 |
| *Daucus carota* | 0,36 | 0,2 | 0,64 | 0,0019 |
| *Diplotaxis erucoides* | 0,29 | 0,79 | 0,48 | 0,0002 |
| *Erigeron canadensis* | 0,79 | 0,29 | 0,55 | 0,0001 |
| *Erodium cicutarium* | 0,47 | 0,29 | 0,43 | 0,0029 |
| *Fumaria officinalis* | 0,08 | 0,43 | 0,39 | 0,0032 |
| *Geranium columbinum* | 0,03 | 0,28 | 0,29 | 0,0019 |
| *Geranium rotundifolium* | 0,28 | 0,47 | 0,51 | 0,0040 |
| *Helminthothec echioides* | 0,49 | 0,16 | 0,66 | 0,0005 |
| *Lactuca serriola* | 0,75 | 0,34 | 0,53 | 0,0005 |
| *Lamium amplexicaule* | 0,86 | 0,23 | 0,47 | 0,0025 |
| *Lolium multiflorum* | 0,19 | 0,3 | 0,5 | 0,0057 |
| *Malva sylvestris* | 0,11 | 0,21 | 0,6 | 0,0017 |
| *Mercurialis annua* | 0 | 0,44 | 0,34 | 0,0036 |
| *Muscari neglectum* | 0,03 | 0,32 | 0,82 | 0,0008 |
| *Plantago lanceolata* | 0,2 | 0,3 | 0,57 | 0,0015 |
| *Poa annua* | 0,28 | 0,54 | 0,68 | 0,0003 |
| *Senecio vulgaris* | 0,43 | 0,72 | 0,47 | 0,0002 |
| *Sonchus asper* | 0,09 | 0,31 | 0,57 | 0,0003 |
| *Sonchus oleraceus* | 0,24 | 0,29 | 0,29 | 0,0003 |
| *Stellaria media* | 0,28 | 0,36 | 0,44 | 0,0004 |
| *Taraxacum officinale* | 0,12 | 0,61 | 0,51 | 0,0007 |
| *Veronica persica* | 0,38 | 0,41 | 0,75 | 0,0010 |
